# Biochemical and Binding Characterization of a Riboflavin Analogue Tethered to Biotin

**DOI:** 10.64898/2026.08.29.748002

**Authors:** Simona Marincean, Sheila R. Smith, Travis Branscum, Amanda Ratajczak, Marilee Benore

## Abstract

The binding affinities of a chimeric analog of a riboflavin derivative linked to biotin, (**6-(7**,**8-dimethyl-2**,**4-dioxo-3**,**4-dihydrobenzo[*g*]pteridin-10(2*H*)-yl)hexyl 5-((3a*S***,**4*S***,**6a*R*)-2-oxohexahydro-1*H*-thieno[3**,**4-*d*]imidazol-4-yl)pentanoate)**, referred to as C6-Rf-biotin-tag, to the riboflavin binding retain or streptavidin are in the μM range, 1.29 ± 0.277 and 3.00 ± 0.459, respectively. These values suggest that C6-Rf-biotin-tag has potential applications in diagnostic assay and labelling target flavin binding proteins. The C6-Rf-biotin-tag which was characterized with respect to physical and biochemical properties retains UV/Vis spectroscopic and fluorescence behavior similar to riboflavin.

**Highlights:**

- A novel molecule of a riboflavin derivative linked to biotin was tested for protein binding
- Fluorescence of the chimeric molecule was quenched upon binding to riboflavin binding protein
- Riboflavin binding protein or streptavidin bind the chimeric molecule
- Plasmon waveguide resonance spectroscopy was used to determine the binding affinities

## 1. Introduction

Metabolic processes are the most fundamental of all physiological reactions; without catabolic energy, organisms die. Riboflavin (Rf), a bright yellow water-soluble small molecule, is a dietary micronutrient that is quickly converted to active coenzymes flavin adenine dinucleotide (FAD) and flavin mononucleotide (FMN) [1]. (Figure 1A). These coenzymes participate in the electron and proton transfers that not only directly impact ATP synthesis but are required in the activation of other cofactors and coenzymes including folate, pyridoxal phosphate, and niacin [2,3]. Rf also plays a critical role in iron metabolism and utilization due to its antioxidant and redox properties [4].

**Figure 1.**
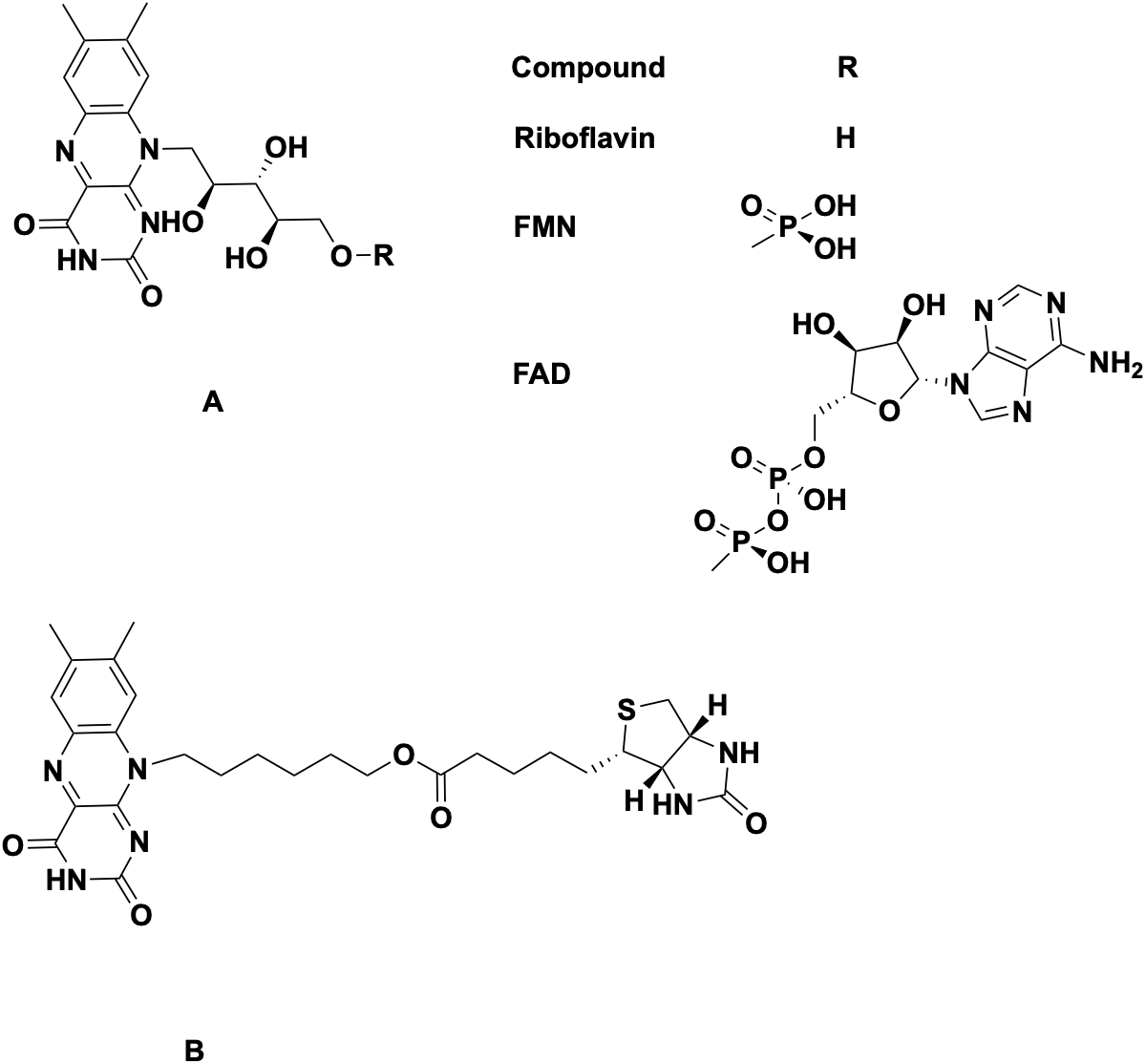
Structures of the flavin coenzymes and the flavin analogue C6-Rf-biotin-tag: A. Rf, FMN and FAD; B. C6-Rf-biotin-tag ((6-(7,8-dimethyl-2,4-dioxo-3,4-dihydrobenzo[g]pteridin-10(2H)-yl)hexyl 5-((3aS,4S,6aR)-2-oxohexahydro-1H-thieno[3,4-d]imidazol

Rf deficiency is common in countries or areas with food shortages, but less so in countries with fortified food policy and nutrient-rich diets. This disparity has led to neglect of Rf research compared to other nutrients [5,6]. Recent studies indicate that flavin deficiency is more widespread than assumed and is found in countries with a range of incomes and resources [7]. Rf deficiency causes or impacts many diseases, and supplementation is often recommended [3]. Ariboflavinosis, depression, and cancer are a few of the diseases linked to flavin status [8,3,9].

Existing methods to measure Rf in vitro or in vivo require expensive instrumentation, assays containing components such as bacteria or red blood cells, or specialized personnel, none easily accessible in remote areas or low-income countries, limiting the ability to develop individual status or a comprehensive picture of global Rf deficiency [10,11]. Thus, better knowledge and factual data of human and animal flavin status and deficiency are important health and wellness goals [12].

Our aim is to create an inexpensive method to detect flavin status by developing an assay to quantitatively measure flavin levels. We recently published the synthesis of a series of Rf analogues, including a chimeric structure in which a Rf derivative is tethered to biotin (**6-(7**,**8-dimethyl-2**,**4-dioxo-3**,**4-dihydrobenzo[*g*]pteridin-10(2*H*)-yl)hexyl 5-((3a*S***,**4*S***,**6a*R*)-2-oxohexahydro-1*H*-thieno[3**,**4-*d*]imidazol-4-yl)pentanoate)**, referred to as C6-Rf-biotin-tag, Figure 1B [13,14].

We describe biochemical characterization of C6-Rf-biotin-tag, including the binding affinity to RBP, a transport protein with high affinity for flavins, and C6-Rf-biotin-tag affinity for avidin and its recombinant form streptavidin [15-18]. These proteins bind their respective natural ligands Rf and biotin with high affinity, and the resulting complexes are used in diagnostic assays and probes [19-23]. C6-Rf-biotin-tag has potential to be used in an assay linking competitive binding of flavin molecules to the biotin/avidin detection system. The physical characteristics of this chimeric molecule were investigated as steps toward developing a series of compounds that can be used in a flavin status detection assay in health applications but also have potential as tools in biochemistry research.

## 2. Materials and Methods

### 2.1 Reagents

Chemical reagents and solvents were purchased (Sigma). The C6-Rf-biotin-tag was synthesized as described, diluted into 100% dimethylsulfoxide (DMSO) and stored in the dark at room temperature [13]. Solutions of Rf or C6-Rf-biotin-tag were further diluted into DMSO or Tris buffer solutions as described. Chicken RBP was purified by the Miller/White method from egg albumen and its purity verified via SDS gel electrophoresis [15,24,25]. Concentrations were determined by spectroscopy using the Kozik equation with a molecular weight of 32,000 D and the extinction coefficients described [26]. RBP was purified from egg white in a mixture containing both bound (holo) and no (apo) Rf ligand. The biotin binding assay was purchased from Sigma. HoloRBP is bright yellow in solid or suspended form, whereas apoRBP is a white solid and transparent in solution. apoRBP was obtained from purified egg white RBP via low pH dialysis to remove the Rf followed by dialysis against H_2_O, and then lyophilization [15]. The removal of Rf was confirmed by spectroscopic analysis; no observed absorbance at 350 or 450 nm regions. Assays for binding were conducted in the buffers as described, typically 10mM Tris pH 8.0 or 10mM Tris pH 8.0/4% DMSO. Rf and C6-Rf-biotin-tag were suspended in water, buffers, or DMSO and diluted as described for the experiments.

### 2.2 Instrumentation

Fluorescence studies were conducted on a Tecan Infinite 200 Pro M Plex microplate reader at excitation and emission wavelengths 450 and 530 nm. UV/Vis spectroscopy experiments were performed on the following instruments: Shimadzu UV-1900I, ThermoFisher Genesys model 140/150, or ThermoFisher NanoDrop One.

### 2.3 Binding of Rf and C6-Rf-biotin-tag to RBP

Binding of C6-Rf-biotin-tag to RBP was demonstrated by fluorescence measurements, as the fluorescence of Rf and other flavin molecules is quenched when bound to RBP [18]. C6-Rf-Biotin-tag or Rf stock was pipetted into microplate wells, increasing amounts of apoRBP added to the wells in distilled water or Tris buffer at room temperature, and fluorescence measured. In addition to the initial readings, absorbance (450 nm) and fluorescence readings were monitored over periods up to 24 hours to ensure binding stability. Control experiments indicated that fluorescence behavior was not affected by the DMSO (data not shown). To test stability, batches of the analogue stored in the dark at room temperature in DMSO for one year were tested to ensure that fluorescence could be quenched by binding to apoRBP (data not shown).

### 2.4 C6-Rf-biotin-tag-RBP complex

It is reported that avidin structural integrity is altered in the presence of DMSO concentrations over 5% DMSO [28]. To enhance the net solubility and minimize concerns about the structural integrity of avidin in the presence of DMSO, a C6-Rf-biotin-tag:RBP complex was prepared: 1 mL 400 μM apoRBP and 2 mL 420 μM C6-Rf-biotin-tag were placed in a 3 mL ThermoFisher Slide-A-Lyzer dialysis cassette, 10K MW cutoff, and dialyzed extensively against successive changes of distilled water to remove unbound ligand. The resulting yellow complex was soluble in water in absence of DMSO and stored frozen. The complex was used in the binding assay studies and other experiments as described and the approximate concentration of C6-Rf-biotin-tag and C6-Rf-biotin-tag:RBP determined as described.

### 2.4 C6Rf-biotin-tag binding to avidin and streptavidin

Biotin and biotin-labeled molecules can be tested for binding to avidin by using the dye reagent 4-Hydroxyazobenzene-2-carboxylic acid (HABA), which binds specifically to avidin but with much lower affinity [29]. Free HABA has a maximum absorbance peak at 350 nm, while bound HABA can be quantified by an absorbance increase at 500 nm. Biotin and biotin labelled molecules will compete with HABA. The Sigma kit H2153, which contains HABA and appropriate buffers, was used for these experiments. The C6-Rf-biotin-tag:RBP complex was used as the competing agent, and controls included apoRBP and buffers. Minor modifications allowed for larger sample volumes while retaining total dilution consistent with kit procedures.

### 2.6 Binding affinity determined by plasmon waveguide resonance (PWR) spectroscopy

The binding affinities of the C6-Rf-biotin-tag for apoRBP and streptavidin were quantified separately via plasmon waveguide resonance (PWR) spectroscopy performed on an Emanant Molecular Interaction Analyzer (Mainline Scientific). Streptavidin was chosen for binding studies because the recombinant avidin demonstrates lower non-specific binding to biotin molecules and is free of biotin; avidin isolated from egg albumen contains some bound biotin. In the case of both proteins, 15 μM C6-Rf-biotin-tag was first immobilized on the surface of a PWR sensor chip that had been pretreated with a (3-aminopropyl)triethoxysilane (APTES) solution. Any portions of the sensor chip surface that did not bind the tag were blocked with bovine serum albumin (BSA), and then the protein of interest (RBP or streptavidin) was introduced and allowed to bind to the immobilized tag until the system reached steady state. This procedure was repeated for five concentrations of each analyte protein, with buffers being matched via buffer exchange as necessary between each step.

## 3.0 Results and Discussion

### 3.1 UV/Vis spectroscopy of C6-Rf-biotin-tag and Rf are similar

UV/Vis Spectroscopy demonstrates that spectra for Rf and C6-Rf-biotin-tag are similar, with UV peaks at 220 and 280 nm, and visible range peaks at ∼365 and 450 nm as shown in Figure 2. Molar extinction coefficients of the C6Rf-biotin-tag were determined to be ε _370_ =7300 M^-^cm^−^ and ε_444_ =9700 M^-^cm^−^ in 10 mM pH 8 Tris buffer with 5%DMSO. These are consistent with literature data on Rf spectrophotometric characteristics, with the yellow isoalloxazine absorbance visible at ∼450 nm. Biotin has a maximum absorbance below 200 nm and does not absorb in the visible region as it has no chromophore. The absorbance peaks of riboflavin shift depending on the pH of the solution.

**Figure 2.**
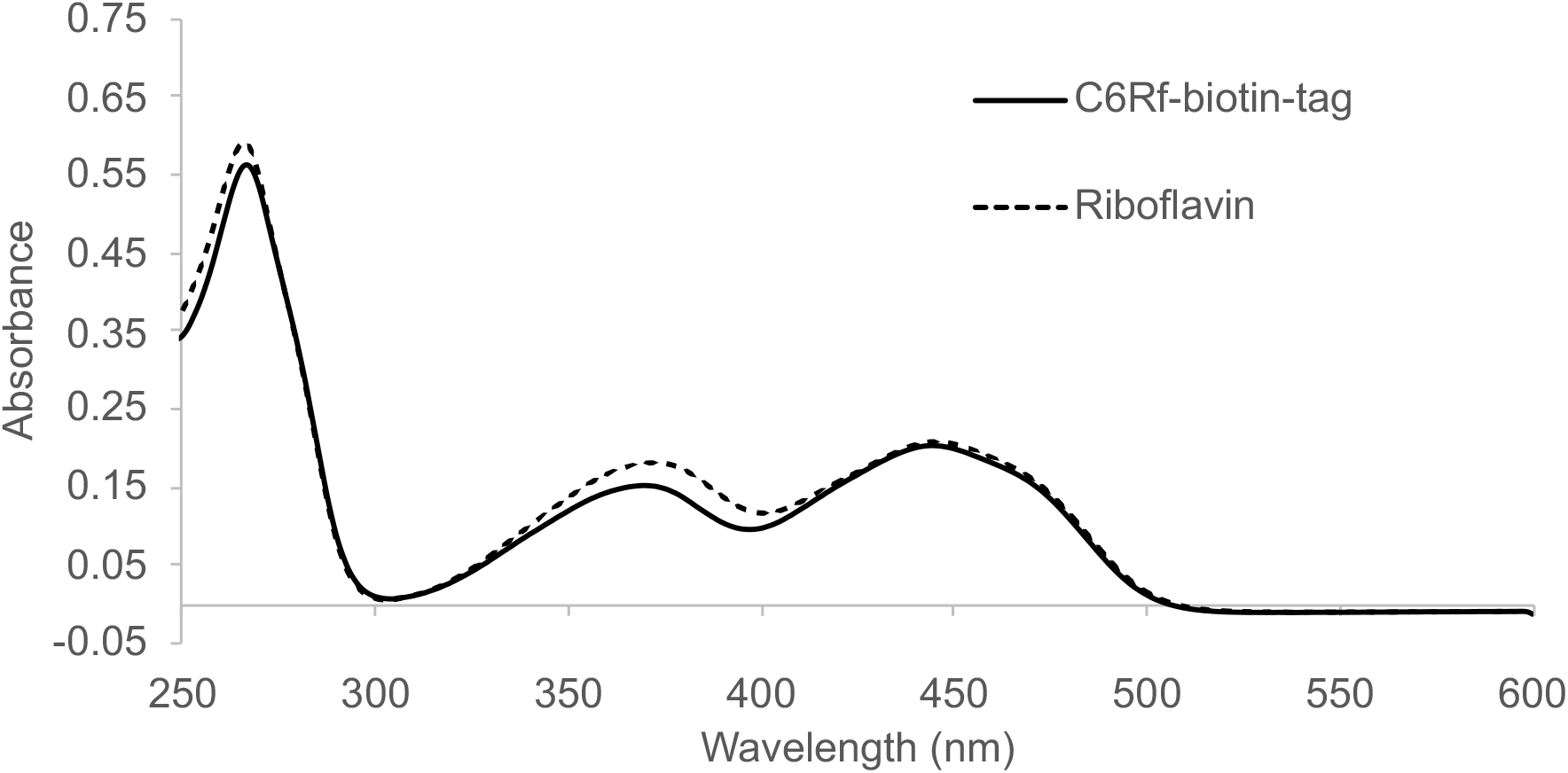
UV/Vis spectroscopy of Rf and C6-Rf-biotin-tag. Each contains 20 □M in 10 mM TRIS pH8 with 5% DMSO

### 3.2 ApoRBP binds both C6-Rf-biotin-tag and Rf

The fluorescence of flavin molecules is quenched upon binding to apoRBP, although the complex is still yellow [10,15,26]. Fluorescence quenching is postulated to be caused by electron sharing between the isoalloxazine ring stacked in the Trp-Tyr binding pocket [30]. Adding increasing amounts of apoRBP to either C6-Rf-biotin-tag or Rf in solution, quenching was observed indicating binding. Fluorescence was measured with emission and excitation wavelengths as described at 450 and 525nm. The data from triplicate samples is shown in Figure 3a. The fluorescence decreases to near zero upon the incremental addition of apoRBP. These experiments were repeated numerous times with similar results. In some experiments the fluorescent measurements were read hourly over a 24-hour period to ensure stability of binding, with little change in the measurements (data not shown). The total fluorescence of the C6-Rf-biotin-tag is significantly lower than that of free Rf, consistent with reports that other flavin analogues, such as FAD, exhibit lower fluorescence.

**Figure 3.**
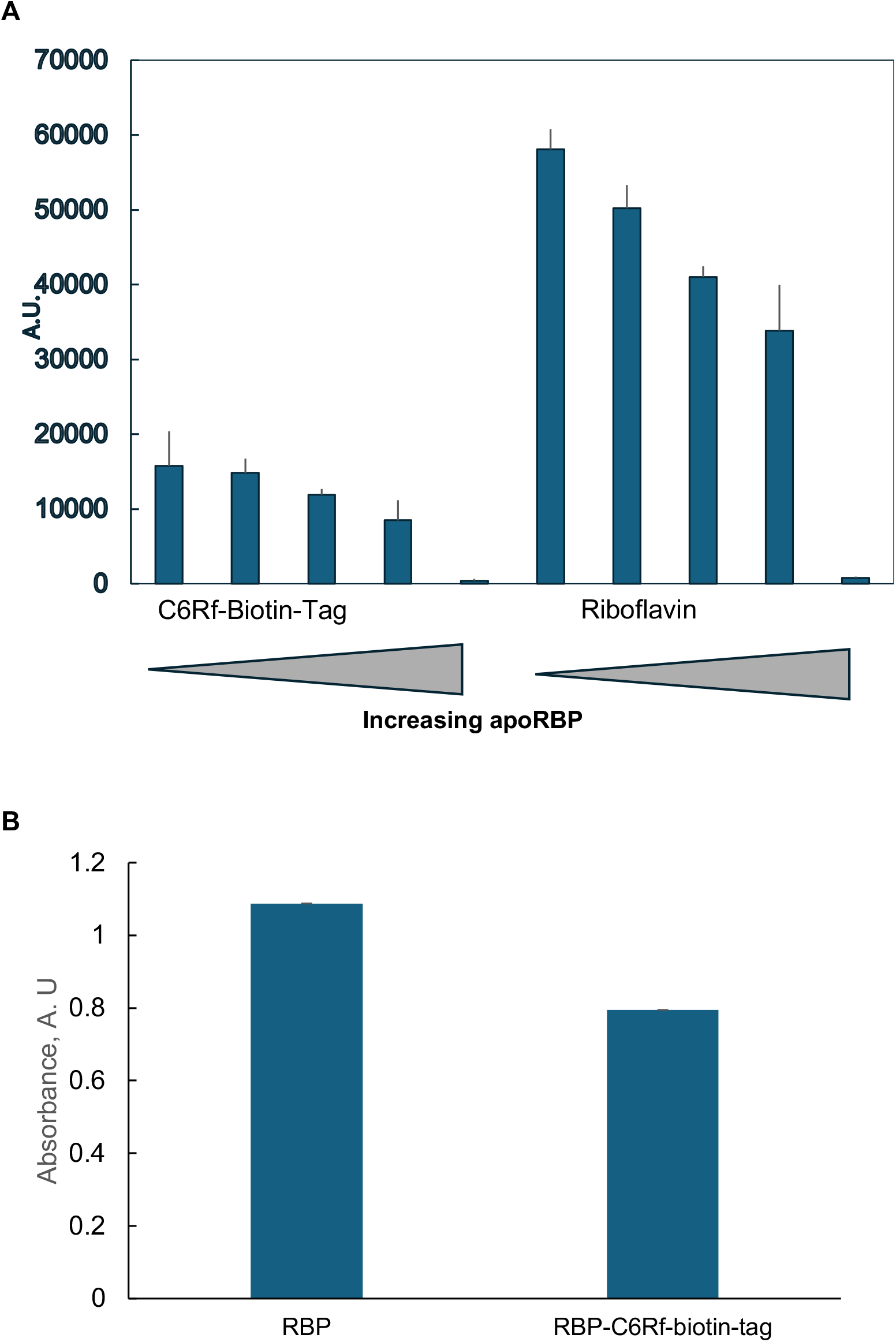
Binding of C6-Rf-biotin-tag to proteins. A. Fluorescence quenching demonstrating binding to RBP, Concentrations: C6-Rf-biotin-tag:16.8 mM; Rf: 23.4 mM; apoRBP: 0, 1.6, 8, 16 and 80 mM; B. Competition with HABA for avidin binding using the Sigma

### 3.3 C6-Rf-biotin-tag binds to avidin

Direct observation of the binding of the C6-Rf-biotin-tag to avidin or streptavidin by fluorescence quenching was not possible in the presence of HABA since HABA also fluoresces. In addition, DMSO present in the ligand samples interfered with the absorbance measurements of the HABA assay. Therefore C6-Rf-biotin-tag:RBP complex was used in the avidin binding experiments. Binding was indirectly measured by competition, as the C6-Rf-biotin-tag displaced the HABA bound to the avidin. The results showed that C6-Rf-biotin-tag added to the HABA-Avidin complex reduced the absorbance at 500nm (Figure 3b). This decrease is within the effective parameters of the Sigma kit. Controls of C6-Rf-biotin-tag were included to ensure that no absorbance change was due to overlap of the isoalloxazine absorbance at 450 nm.

### 3.4 Binding Affinities of C6-Rf-biotin-tag for RBP and streptavidin

The PWR steady state binding assay results were plotted as shown in Figure 4. Analysis revealed K_D_ values of (3.00 ± 0.459) μM and (1.29 ± 0.277) μM for the binding of streptavidin and apoRBP, respectively, to the C6-Rf-biotin-tag.

**Figure 4.**
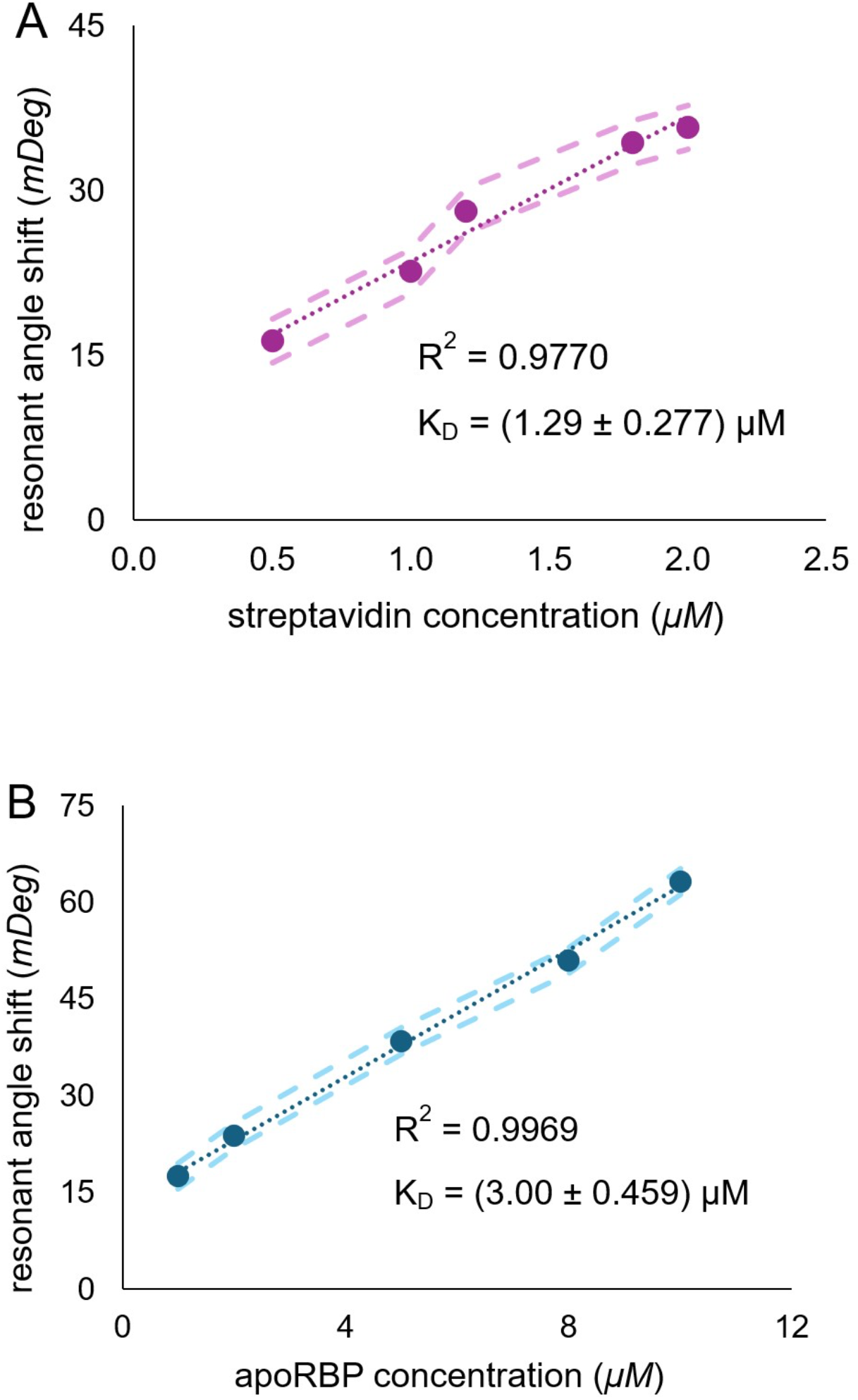
Binding Affinity Determination. A. C6-Rf-biotin-tag at increasing concentrations of Streptavidin; B. C6-Rf-biotin-tag binding at increasing concentrations of RBP.

## Conclusion

The C6-Rf-biotin-tag, with a chimeric structure containing both biotin and modified Rf moieties, binds to both RBP and avidin. We have demonstrated by direct and competition studies that the binding is specific and stable. Affinity was determined as (3.00 ± 0.459) μM for C6-Rf-biotin-tag binding to Streptavidin and (1.29 ± 0.277) μM to RBP. These are lower than the published affinities of biotin for avidin, at 1.3 x 10^-15^ M, and Riboflavin binding to RBP in the range of 1-40 nM. The affinities are consistent with the lower values observed of other ligands binding to RBP or avidin [18, 29].

We were unable to demonstrate simultaneous binding of C6-Rf-biotin-tag to RBP and avidin. If the complex C6-Rf-biotin-tag-RBP was able to compete with the HABA, we would expect fluorescence quenching to be unchanged, but if the C6-Rf-Biotin-Tag was free, fluorescence would increase upon removal form RBP and binding to streptavidin. Unfortunately, the inherent fluorescence of the HABA-Avidin reaction occurs at similar wavelengths, making it impossible to determine if quenching was changed. We suspect that the very high affinity of the avidin for biotin attracted the C6-Rf-biotin-tag, detaching it from the RBP.

This is not unexpected as the binding sites of the individual moieties are well known and we believe simultaneous binding unlikely due to steric strain. RBP binds Rf with the isoalloxazine buried in a cleft flanked by Trp and Tyr residues, with a small portion of the tail on the surface. [30] Biotin binds to a groove on avidin, with little of the tail exposed [31]. (Protein data base PDB entry 2AVI) Thus, the proteins’ bulk at the binding sites’ surfaces would overlap, resulting in steric hindrance.

A chimeric molecule of Rf linked to biotin has potential for quantitative diagnostic assay development, as a marker for receptor research and diagnosis, or as an inhibitor for bacterial studies. Future work on will steps include synthesis of chimeric molecules with longer linking tethers or altered isoalloxazines to increase solubility, increase binding affinity and augment simultaneous binding to RBP and avidin.

## Contributions

**Simona Marincean:** Conceptualization, Funding acquisition, Investigation, Methodology, Project administration, Writing – review and editing: **Sheila R. Smith:** Investigation, Methodology, Project administration, Writing – review and editing: **Travis Branscum** : Investigation: **Amanda Ratajczak**: Investigation, Methodology, Writing – review and editing: **Marilee Benore:** Conceptualization, Funding acquisition, Investigation, Methodology, Project administration, Writing – original draft, Writing – review and editing **Funding:** This research received no external funding

## Acknowledgments

We acknowledge financial support from the University of Michigan-Dearborn Office of Sponsored Research (ORSP). We thank Larry Medina of Mainline Scientific for helpful discussions.

## Declaration of competing interests

The authors declare no conflicts of interest but have submitted a patent: United States patent application # PCT/US25/41035

## Abbreviations

The following abbreviations are used in this manuscript

I: current (A)
Rf: Riboflavin
ApoRBP: Apo Riboflavin Binding Protein DMSO Dimethyl Sulfoxide
Tris: Tris(hydroxymethyl)aminomethane
FAD: Flavin adenine dinucleotide
FMN: Flavin mononucleotide
C6Rf-biotin-tag: (6-(7,8-dimethyl-2,4-dioxo-3,4-dihydrobenzo[*g*]pteridin-10(2*H*)-yl)hexyl 5-((3a*S*,4*S*,6a*R*)-2-oxohexahydro-1*H*-thieno[3,4-*d*]imidazol-4-yl)pentanoate),

## Notes

### Competing Interest Statement

The authors have declared no competing interest.

